# Integration of probabilistic functional networks without an external Gold Standard

**DOI:** 10.1101/2021.10.01.462727

**Authors:** Katherine James, Aoesha Alsobhe, Simon J. Cockell, Anil Wipat, Matthew Pocock

## Abstract

**Background:** Probabilistic functional integrated networks (PFINs) are designed to aid our understanding of cellular biology and can be used to generate testable hypotheses about protein function. PFINs are generally created by scoring the quality of interaction datasets against a Gold Standard dataset, usually chosen from a separate high-quality data source, prior to their integration. Use of an external Gold Standard has several drawbacks, including data redundancy, data loss and the need for identifier mapping, which can complicate the network build and impact on PFIN performance.

**Results:** We describe the development of an integration technique, ssNet, that scores and integrates both high-throughput and low-throughout data from a single source database in a consistent manner without the need for an external Gold Standard dataset. Using data from *Saccharomyces cerevisiae* we show that ssNet is easier and faster, overcoming the challenges of data redundancy, Gold Standard bias and ID mapping, while producing comparable performance. In addition ssNet results in less loss of data and produces a more complete network.

**Conclusions:** The ssNet method allows PFINs to be built successfully from a single database, while producing comparable network performance to networks scored using an external Gold Standard source.

## Background

Probabilistic functional integrated networks (PFINs) combine functional interaction data to produce a network of interactions that are annotated with confidence scores. These scores are often calculated by comparison with a high quality “Gold Standard” dataset [1–5]. The performance of any PFIN is dependent upon its component datasets and on the Gold Standard(s) chosen to evaluate them [6]. Integrated network theory relies on the principle that the whole network is greater than the sum of its dataset parts, so as more interaction data are produced, network performance should increase [7]; a PFIN integrated using data from 2010 is expected to perform better than one using data from 2006. However, our previous 2012 study, which investigated PFINs integrated using data from 2006 to 2010, demonstrated that the network performance fluctuated and that careful selection of data and Gold Standard is vital [8], particularly if the network is integrated to study a specific area of biology.

Since 2012 the volume of functional interaction data has increased considerably [9]. While high-throughput (HTP) data produced from largescale screening have been estimated to have high false positive and false negative rates (up to 91% and 96%, respectively) [1, 10–15], the effect of this noise should be mitigated by an increase in less error-prone targeted low-throughput (LTP) studies. Curated databases are constantly evolving in both content and structure to reflect current biological knowledge [16]. Manual curation has improved the quality of available data by editing or removal when errors and inconsistencies have been identified and when data are found to be incorrect in light of new studies [17, 18].

Gold Standard data are often chosen from a separate high-quality database, commonly manually-curated metabolic pathways [1, 19, 20], for example KEGG [21], or shared biological process [10, 22, 23], for example Gene Ontology [24]. The difference in database source between the datasets to be scored and Gold Standard presents several challenges to scoring and integration. The Gold Standard may have poor overlap with the datasets since its focus differs, for instance a metabolic pathway database to score protein-protein interaction (PPI) data; many PPI datasets may not score at all. Conversely, data redundancy can occur since separate databases may include the same studies, leading to overlap between Gold Standard and datasets, and biased scoring. Identifier format may also differ, requiring potentially error prone mapping between the two sources prior to scoring [25]. In Gold Standards derived from hierarchical annotation schemas, such as Gene Ontology, the choice of which annotation terms to include in the Gold Standard can vastly change the final scores and these schemas are known to have large biases [26].

With the increase in volume of high quality LTP data, we hypothesise that these data can be used as a Gold Standard to score the larger HTP studies. Therefore, using functional interaction data from a single curated database, datasets can be scored and integrated without the requirement for an external Gold Standard dataset. Less data will be lost and the challenges of identifier mapping and data redundancy will be removed. Furthermore, since the Gold Standard is considered the highest-quality data, we propose that these data may be integrated into the final network using iterative scoring.

Here, we extend our 2012 study by analysing the changes in *Saccharomyces cerevisiae* functional interaction data contained in one of the most comprehensive manually-curated databases, BioGRID^1^ [27]. We find that the quantity of interaction data has increased to the extent whereby both scoring and integration can be achieved from the data contained in BioGRID alone. We propose a new integration method, ssNET, that incorporates both LTP and HTP data in a consistent manner, and evaluate it using functional prediction.

The networks produced in this study and the code for ssNet network generation are available at: https://figshare.com/projects/Yeast_ssNet/114366 and https://github.com/kj-intbio/ssnet/.

## Results

### LTP data can be used as a Gold Standard to score HTP data quality

PFINs are traditionally integrated after datasets have been scored against an external Gold Standard dataset [1, 19, 20, 22]. Since the use of external data has several drawbacks, we investigated whether the less error-prone low-throughput (LTP) studies are suitable to be used as an internal Gold Standard. We first assessed the changes in BioGRID data for the yeast *Saccharomyces cerevisiae* from the first available version (V17, 2006) to the current version (V186, 2020). HTP interaction data (defined here as studies *>*= 100 interactions; see Methods) have increased considerably since our earlier study [8] both in terms of unique proteins and in unique interactions between them (Figure 1 A and B). The overlap between HTP and LTP interactions remains relatively low, increasing from 3382 (7.1% of total interactions) at V17 to 16724 interactions at V186 (3.1% of total interactions). While the number of unique proteins fluctuated in earlier versions of BioGRID, their numbers have steadily increased in later years. The proportion of protein overlap between HTP and LTP data has also increased from 3377 at V17 to 5016 at V186: 63.8% and 81.5% of total proteins, respectively. Increasing the size threshold for the LTP datasets from 100 interactions had little effect on the percentage overlap between HTP and LTP datasets (Figure 1 C and D).

**Figure 1:**
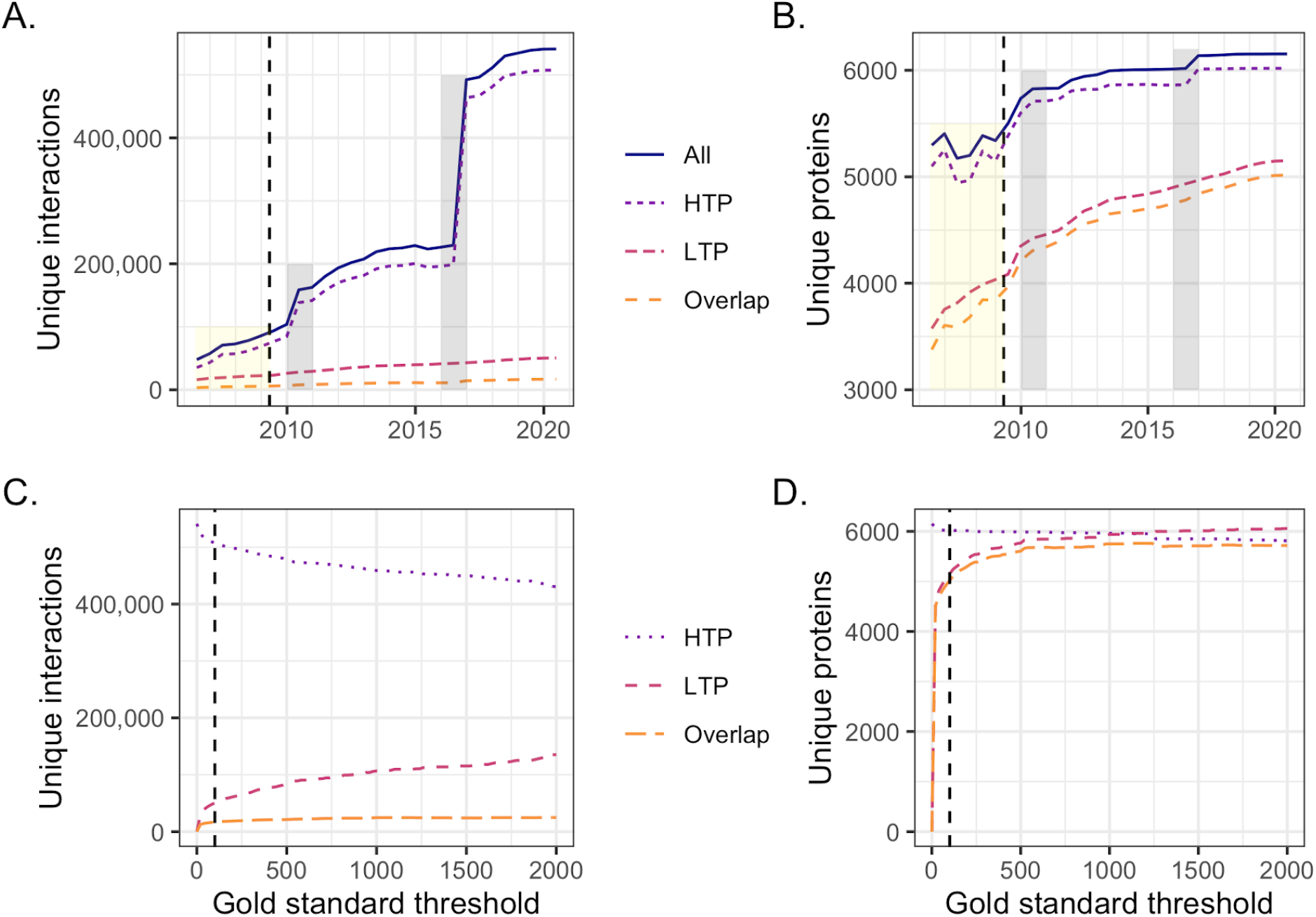
The BioGRID *Saccharomyces cerevisiae* data increase. A. High-throughput (HTP) interaction data has increased since our earlier study [8] (yellow box; dashed line), with two large increases in 2010 and 2016 (grey boxes). By contrast, low-throughput (LTP) data has increased to a lesser extent, with the overlap between HTP and LTP also increasing slightly. B. The fluctuation in representation of unique proteins that was observed in our earlier study [8] (yellow box; dashed line) is not evident in the following years. Unique protein number has increased steadily for both HTP and LTP data and the large increases in interaction data (grey boxes) only affect this increase slightly. Version 186 LTP data contains ∼ 1*/*3 of the total unique proteins in BioGRID, with LTP data almost completely overlapping with the HTP dataset. C. The size and overlap of interactions in V186 HTP datasets and LTP Gold Standard data as the Gold Standard size threshold is increased. D. The size and overlap of proteins in V186 HTP datasets and LTP Gold Standard data as the Gold Standard size threshold is increased. An LTP dataset size threshold of 100 interactions (dashed line) was chosen for subsequent analyses to maximise both the number of the LTP Gold Standard’s unique proteins and unique interactions, while minimising its overall size.

We constructed an LTP-derived Gold Standard (LTP GS) and three Gold Standards derived from the metabolic pathways of the BioSystems database [28] (MP GS) and biological processes of the Gene Ontology (BP10 GS and BP100 GS) [24] for comparison (see Methods). An LTP dataset size threshold of < 100 interactions was chosen to maximise the number of unique proteins and unique interactions in the LTP GS, while minimising the its overall size (Figure 1 C and D).

The LTP GS had greater overlap with HTP data in terms of individual proteins, with ∼ 83% of the total proteins shared (Table 1), in comparison to ∼ 42% for MP GS, ∼ 81% for BP10 GS and ∼ 73% for BP100 GS. These results suggest that while overlap between LTP and HTP data is low, LTP data has better overlap than the external Gold Standard in terms of proteins, and therefore may provide improved scoring. The LTP GS also had higher overlap with the HTP datasets than MP GS in terms of interactions with 3.3% in comparison to 0.2%, but lower overlap than BP10 GS (34.5%) and BP100 GS (7.73%). While the largest Gold Standard, BP10 GS has the highest overlap with the HTP datasets, it’s size may adversely affect scoring since the prior expectation (*P*(*L*); see Methods) will contain a large number of linkages between proteins with shared biological process that are less likely to interact directly.

**Table 1:** Gold Standard overlap. The overlap in terms of proteins and interactions of the high-throughput datasets (HTP) with the low-throughout Gold Standard (LTP_GS), the BioSystems-derived Gold Standard (MP_GS) and Gene Ontology biological process-derived Gold Standards (BP10_GS and BP100_GS) (highest % overlaps are shown in bold). Total numbers differ slightly from Figure 1 since only those identifiers that could be mapped between Gold Standard and BioGrid are included.

|  |  | LTP_GS | MP_GS | BP10_GS | BP100_GS |
| --- | --- | --- | --- | --- | --- |
| **Proteins**<br>(6,018 HTP) | Gold Standard | 5,510 | 2,551 | 5,071 | 4,551 |
|  | Overlap | 5,016 | 2,549 | 4,868 | 4,369 |
|  | Overlap % | **83.3** | 42.4 | 80.9 | 72.6 |
| **Interactions**<br>(507,352 HTP) | Gold Standard | 50,491 | 9,924 | 2,947,156 | 384,450 |
|  | Overlap | 16,724 | 1,045 | 174,934 | 39,200 |
|  | Overlap % | 3.3 | 0.2 | **34.5** | 7.73 |

When these Gold Standards were used in dataset scoring the majority of scores were higher using the LTP_GS in all three cases. The scores also had greater spread, indicating that the LTP_GS provides wider scope to score more diverse datasets (Figure 2). Datasets may score infinity if they have perfect overlap with the Gold Standard (¬*P*(*L*|*E*) = 0, see Methods). No datasets scored infinity using the LTP_GS, in comparison to 10 datasets using MP_GS, 2 using BP10_GS and 1 using BP100_GS (shown in Figure 2 as a score of 7.0 on the right of the plot).

**Figure 2:**
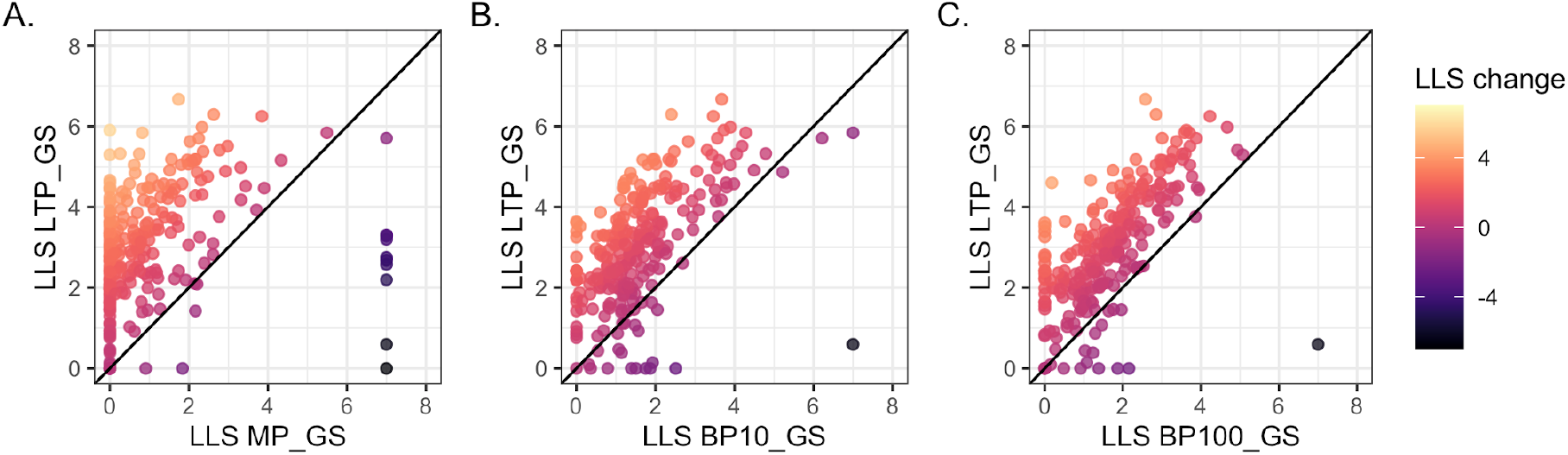
Dataset scoring. The log likelihood scores (LLS) comparison for V186 BioGRID datasets scored against the low throughput (LTP_GS). Datasets that had perfect overlap with Gold Standard (¬*P*(*L*|*E*) = 0) so are shown as score 7.0. Datasets shown as 0 were lost due scoring <0 or no score (*P*(*L*|*E*) = 0). A. BioSystems-derived (MP GS) Gold Standard (5 datasets score 0 for both Gold Standards). B. Gene ontology biological process terms annotating <10% of the genome (BP10 GS) Gold Standard. C. Gene ontology biological process terms annotating annotation <100 genes (BP100_GS) Gold Standard (2 datasets score 0 for both Gold Standards).

Datasets may also be lost during scoring if they do not have any overlap with the Gold Standard data (*P* (*L* |*E*) = 0, see Methods), or have negative scores. Fewer datasets were lost when scored against the LTP_GS (8 with LTP_GS, 96 with MP_GS, 18 with BP10 GS and 23 with BP10 GS; shown as score 0 in Figure 2). Since the most recent BioSystems dataset available was April 2017, we also confirmed the difference between LTP_GS and MP_GS using equivalent BioGRID data from 2017 (V150, data not shown). The use of LTP data as Gold Standard for scoring therefore results in less loss of data than using the external Gold Standards.

Four networks were integrated using V186 HTP datasets scored against the LTP_GS and the MP_GS, BP10 GS and BP100_GS Gold Standards for comparison. The networks were all formed of one connected component, with the LTP_GS network being larger and having the largest average degree (168 compared to 155 for MP_GS, 167 for BP10 GS and 166 for BP100_GS) (Table 2). The networks all had moderate correlation with the power law [29], possibly a reflection of their content being solely derived from noisy HTP data. The high connectivity of the LTP_GS network was reflected in the clustering output with the network being split into 87 clusters, compared to 220 clusters for the MP_GS-scored network and 169 for BP100_GS. BP10_GS gave the fewest clusters with just 31.

**Table 2:** Network topology. Network statistics for the low throughput (LTP_GS), BioSystems (MP_GS) and Gene Ontology (BP10_GS and BP100_GS) scored networks. Statistics were calculated using the Cyctoscape NetworkAnalyser [30] plugin and clustering was carried out using the Markov Clustering Algorithm (MCL) [31].

|  | **LTP_GS** | **MP_GS** | **BP10_GS** | **BP100_GS** |
| --- | --- | --- | --- | --- |
| Proteins | 5,960 | 5,919 | 5,960 | 5,958 |
| Interactions | 501,299 | 458,778 | 498,385 | 495,007 |
| Average degree | 168 | 155 | 167 | 166 |
| Connected components | 1 | 1 | 1 | 1 |
| Diameter | 4 | 5 | 4 | 4 |
| Characteristic path length | 2.07 | 2.15 | 2.07 | 2.07 |
| Centralization | 0.57 | 0.33 | 0.58 | 0.57 |
| Correlation Power law | 0.41 | 0.46 | 0.41 | 0.43 |
| $R^2$ Power law | 0.74 | 0.75 | 0.74 | 0.74 |
| Clusters | 87 | 220 | 31 | 169 |

We compared the performance of the networks in prediction of 398 Gene Ontology (GO) biological process terms using leave one out prediction of known annotations measured using area under curve (AUC) of receiver operator characteristic (ROC) plots. The LTP_GS network had improved performance for the majority of GO terms in comparison to MP_GS (Figure 3 A and B) with of terms with 327 improved compared to 71. Of the AUC changes 227 were statistically significant with 209 (92%) improved using LTP_GS. Performance was also improved over BP10_GS with 230 terms improved compared to 168. Of the changes 114 were significant with 64 (56%) improved using LTP_GS. LTP_GS and BP100_GS had the most similar performance with 105 terms improved for LTP_GS compared to 293. However, only 85 changes were statistically significant and with just 3 (4%) were improved using LTP_GS.

**Figure 3:**
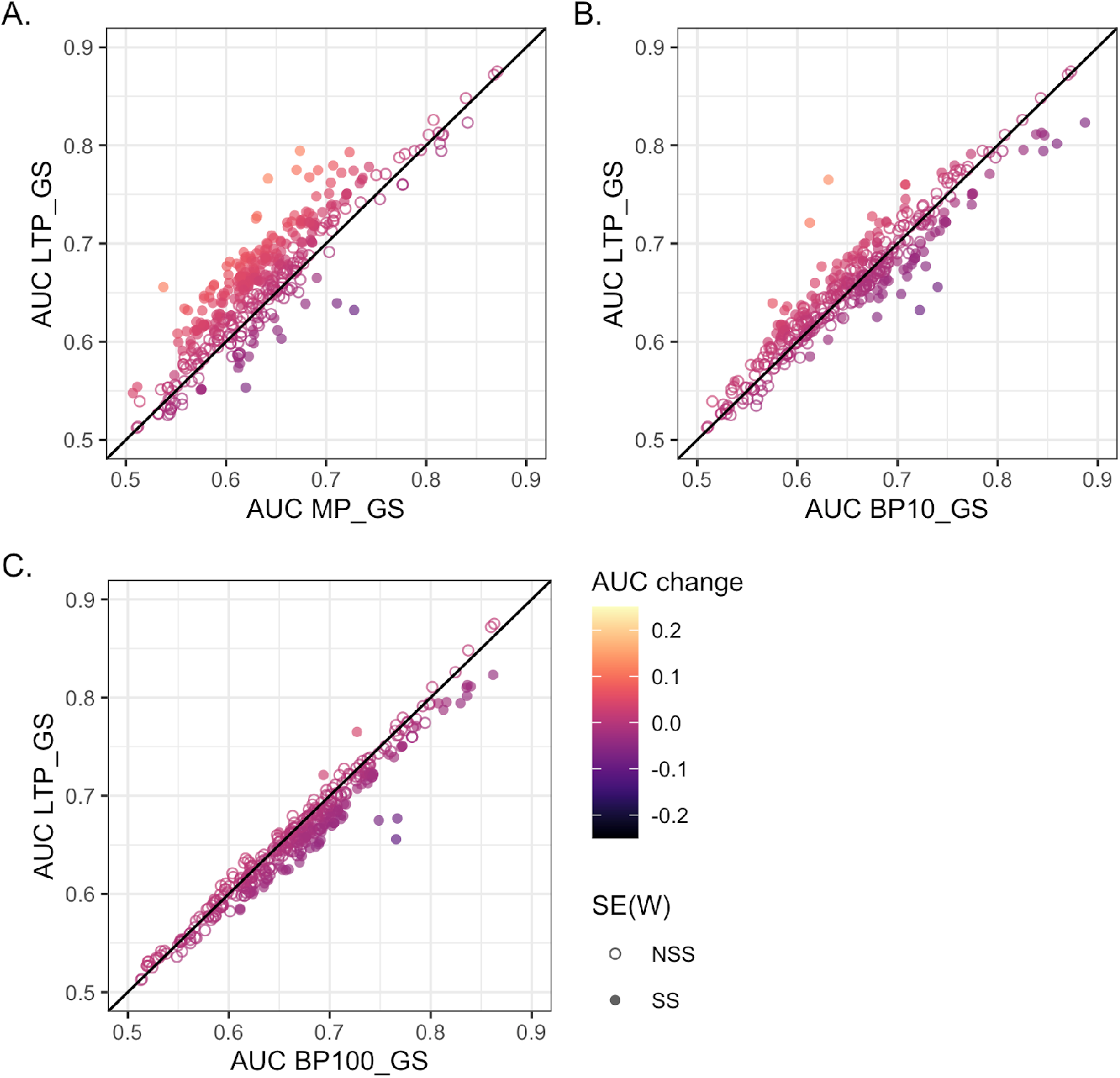
Network evaluation. The functional prediction performance of the low throughput (LTP_GS) and BioSystems and Gene Ontology-scored networks as measured by area under curve (AUC) of receiver operator characteristic plots. The error of the AUC was calculated using the standard error of the Wilcoxon statistic SE(W): not statistically significant (NSS); statistically significant (SS). A. BioSystems-derived (MP_GS) Gold Standard. Of 398 Gene Ontology biological processes, 327 had improved prediction using the LTP_GS network, and 71 using the MP_GS network. Of these changes 227 were statistically significant with 209 (92%) improved using LTP_GS. B. Gene ontology biological process terms annotating <10% of the genome (BP10_GS) Gold Standard. Of 398 Gene Ontology biological processes, 230 had improved prediction using the LTP_GS network, compared to 168 using BP10_GS. Of the changes 114 were significant with 64 (56%) improved using LTP_GS. C. Gene ontology biological process terms annotating annotation <100 genes (BP100_GS) Gold Standard. Of 398 Gene Ontology biological processes, 105 had improved prediction using the LTP_GS network, compared to 293 using BP100_GS. Of the changes just 85 were significant with 3 (4%) improved using LTP_GS.

### Incorporation of Gold Standard data improves network performance

Since the LTP_GS data are considered the highest-quality, their inclusion in the final network is desirable. We therefore investigated how LTP data could be scored and incorporated into the final integrated network. Since LTP data can be further divided by experimental type [19], we employed an iterative LTP scoring method, ssNet, in which each LTP data type is scored as a single dataset using the remaining LTP types as Gold Standard, before integration of the LTP and HTP dataset scores (see Methods). Scores for the 28 LTP datasets were in a higher but overlapping range to those of the HTP datasets using ssNet (Figure 4 A), while LTP scores using the other Gold Standards did not reflect the higher quality of these datasets (Figure 4 B-D). LTP data scores where higher in comparison to scores using a the other three Gold Standards (Figure 5; five LTP datasets that did not score using MP_GS are shown as score 0).

**Figure 4:**
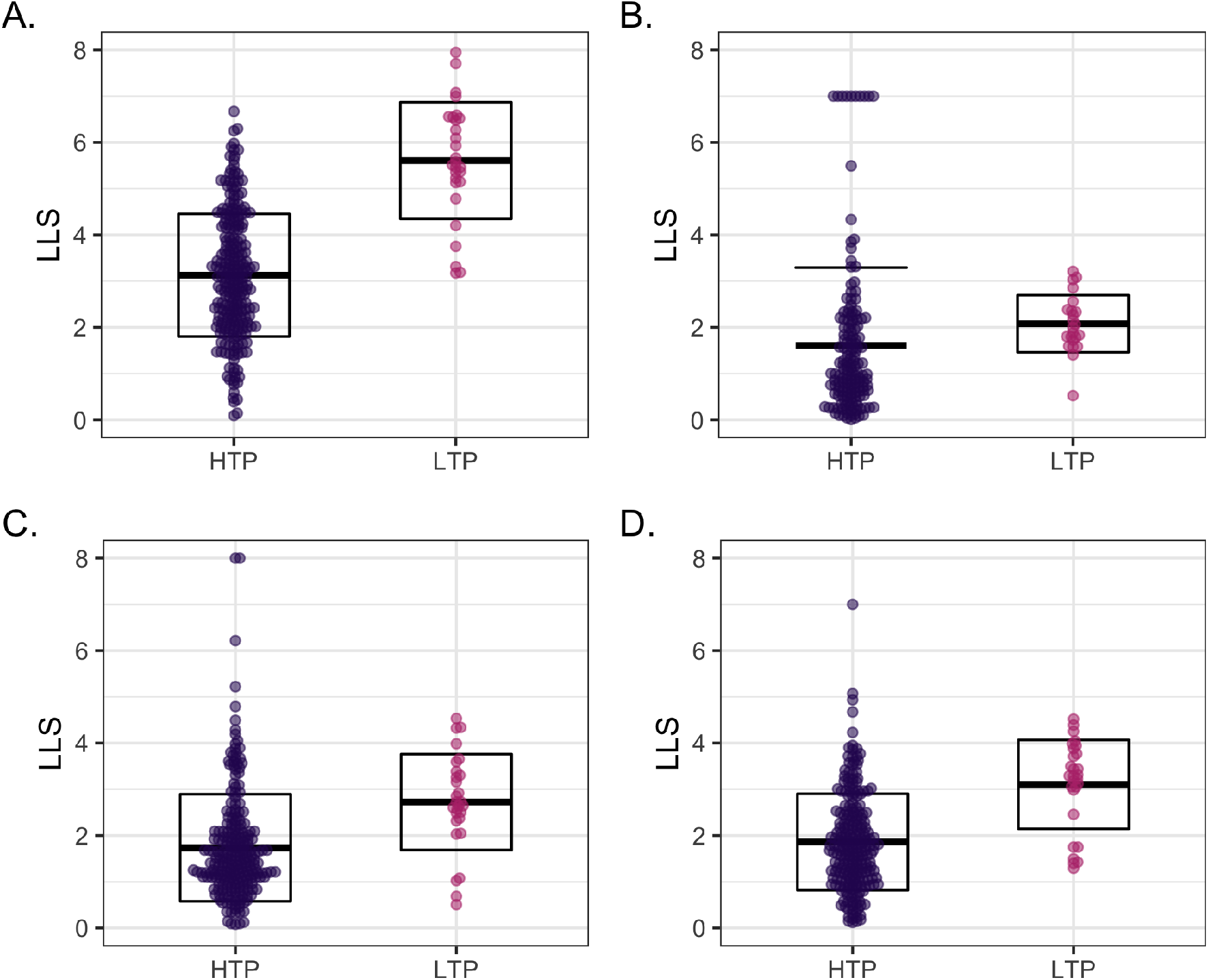
LTP and HTP scores. A. Comparison of log-likelihood (LLS) score range for low-throughput (LTP) and high-throughput (HTP) datasets scored using ssNet. B. Comparison of log-likelihood (LLS) score range for low-throughput (LTP) and high-throughput (HTP) datasets scored using MP_GS. C. Comparison of log-likelihood (LLS) score range for low-throughput (LTP) and high-throughput (HTP) datasets scored using BP10_GS. D. Comparison of log-likelihood (LLS) score range for low-throughput (LTP) and high-throughput (HTP) datasets scored using BP100_GS.

**Figure 5:**
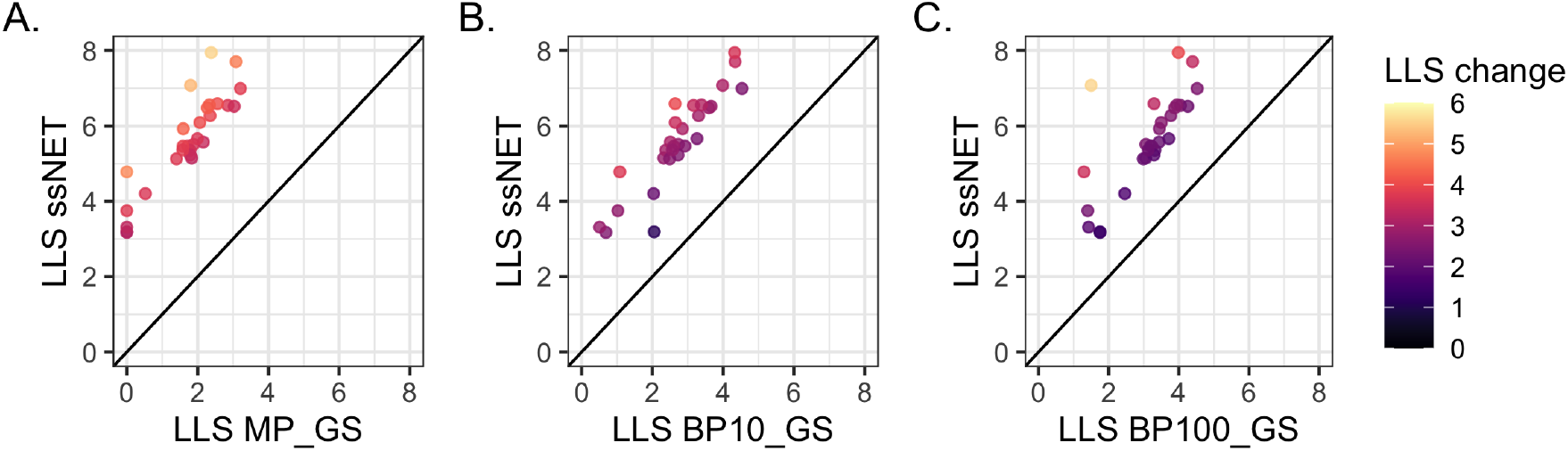
ssNet scoring. A. Comparison of LTP dataset scores using ssNet and the BioSystems-derived Gold Standard (MP_GS). Datasets which were lost due scoring <0 or no score (*P*(*L*|*E*) = 0) are shown as 0. B. Comparison of LTP dataset scores using ssNet and the Gene Ontology Gold Standard BP10_GS. C. Comparison of LTP dataset scores using ssNet and the Gene Ontology Gold Standard BP100_GS.

We integrated four networks of V186 LTP and HTP data scored using ssNet and MP_GS, BP10_GS and BP100_GS Gold Standards, for comparison, and assessed their performance at prediction of protein function as before. The ssNet, BP10_GS and BP100_GS networks were formed of one connected component, while MP_GS had a second small component consisting of three proteins. The ssNet network was larger and more tightly connected (Table 3) with >535,000 interactions and all networks had higher correlation with the power law than those based on HTP data alone (Table 2). We compared the performance of the networks in prediction of 398 Gene Ontology (GO) biological process terms as before. The results were similar to the HTP-only networks with the ssNet having improved performance for the majority of GO terms in comparison to MP_GS (Figure 6 A and B) with of terms with 325 improved compared to 73. Of the AUC changes 127 were statistically significant with 124 (98%) improved using LTP_GS. Performance was not improved over BP10_GS with 128 terms improved compared to 270. Of the changes just 20 were significant with 2 (10%) improved using LTP_GS. LTP_GS and BP100_GS had the most similar performance with 102 terms improved for LTP_GS compared to 296. However, only 2 changes were statistically significant both of which were improved using BP100_GS.

**Table 3:** Network statistics. Topological characteristics for the ssNet, BioSystems (MP_GS) and Gene Ontology (BP10_GS and BP100_GS) scored networks (including the LTP datasets). Statistics are calculated for the larger (6036 proteins) component of the BioSystems network.

|  | ssNet | MP_GS | BP10_GS | BP100_GS |
| --- | --- | --- | --- | --- |
| Proteins | 6,119 | 6,039 | 6,119 | 6,118 |
| Interactions | 535,183 | 492,424 | 532,349 | 529,036 |
| Average degree | 175 | 163 | 174 | 173 |
| Connected components | 1 | 2 | 1 | 1 |
| Diameter | 6 | 5 | 6 | 6 |
| Characteristic path length | 2.11 | 2.16 | 2.11 | 2.11 |
| Centralization | 0.56 | 0.33 | 0.56 | 0.56 |
| Correlation Power law | 0.61 | 0.58 | 0.60 | 0.61 |
| R<sup>2</sup> Power law | 0.74 | 0.76 | 0.74 | 0.74 |
| Clusters | 308 | 394 | 263 | 359 |

**Figure 6:**
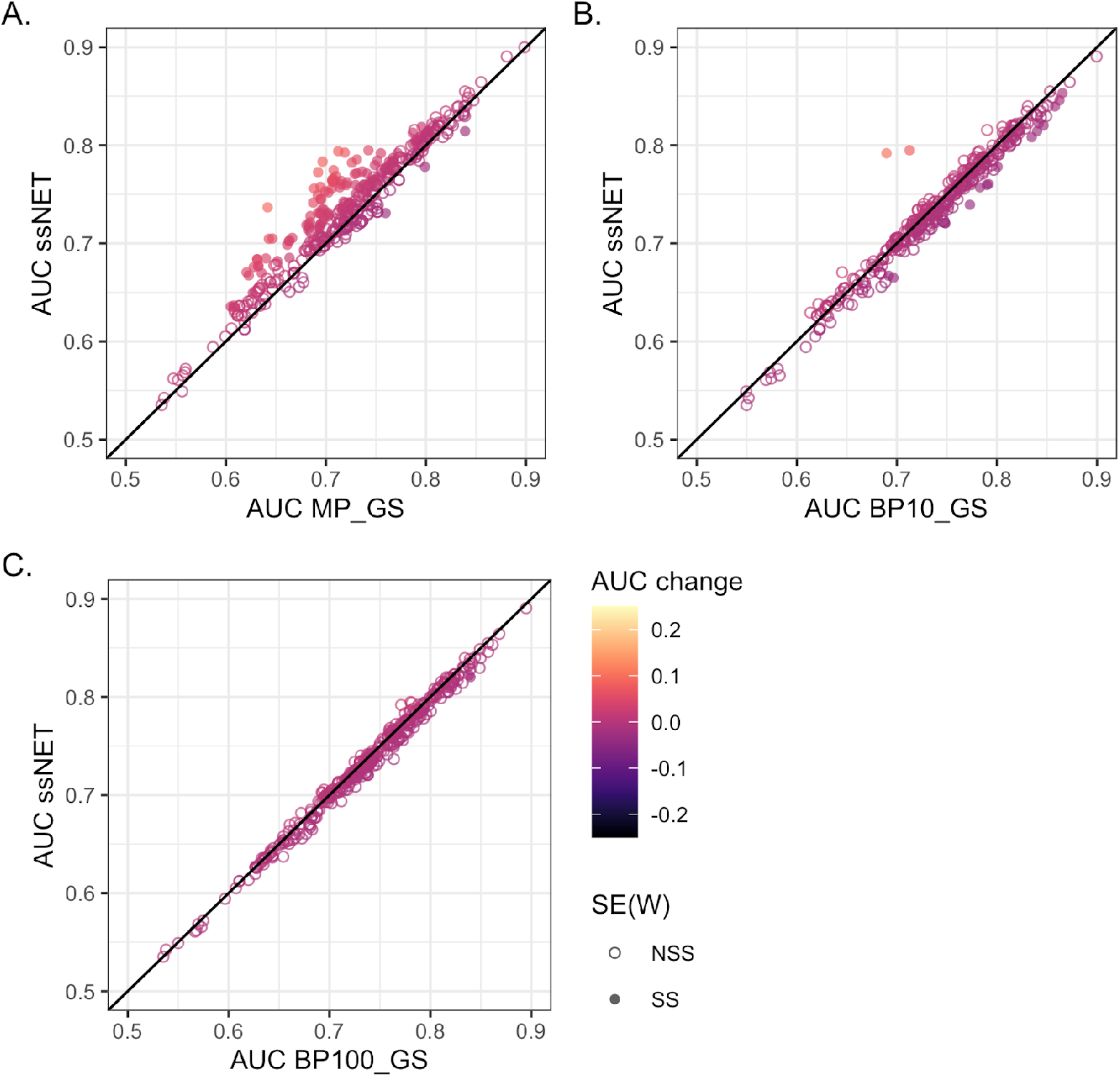
ssNet evaluation. The functional prediction performance of ssNet and BioSystems and Gene Ontology-scored networks as measured by area under curve of AUC of receiver operator characteristic (ROC) plots. The error of the AUC was calculated using the standard error of the Wilcoxon statistic SE(W): not statistically significant (NSS); statistically significant (SS). A. BioSystems-derived (MP_GS) Gold Standard. Of 298 Gene Ontology biological processes, 325 had improved prediction using the ssNet network, and 73 using the MP_GS network. Of these changes 127 were statistically significant with 124 (98%) improved using LTP_GS. B. Gene ontology biological process terms annotating <10% of the genome (BP10_GS) Gold Standard. Of 398 Gene Ontology biological processes, 128 had improved prediction using the LTP_GS network, compared to 270 using BP10_GS. Of the changes just 20 were significant with 2 (10%) improved using LTP_GS. C. Gene ontology biological process terms annotating annotation <100 genes (BP100_GS) Gold Standard. Of 398 Gene Ontology biological processes, 102 had improved prediction using the LTP_GS network, compared to 296 using BP100_GS. Of the changes only 2 were significant, both of which were improved using BP100_GS.

Finally, we investigated the ssNet network’s performance when integrated using different *D* -values (see Methods). In general networks performance is better at lower *D* -values (Figure 7 A). However, individual Gene Ontology biological processes have variation in performance with differing optimal *D* -values (Figure 7 B).

**Figure 7:**
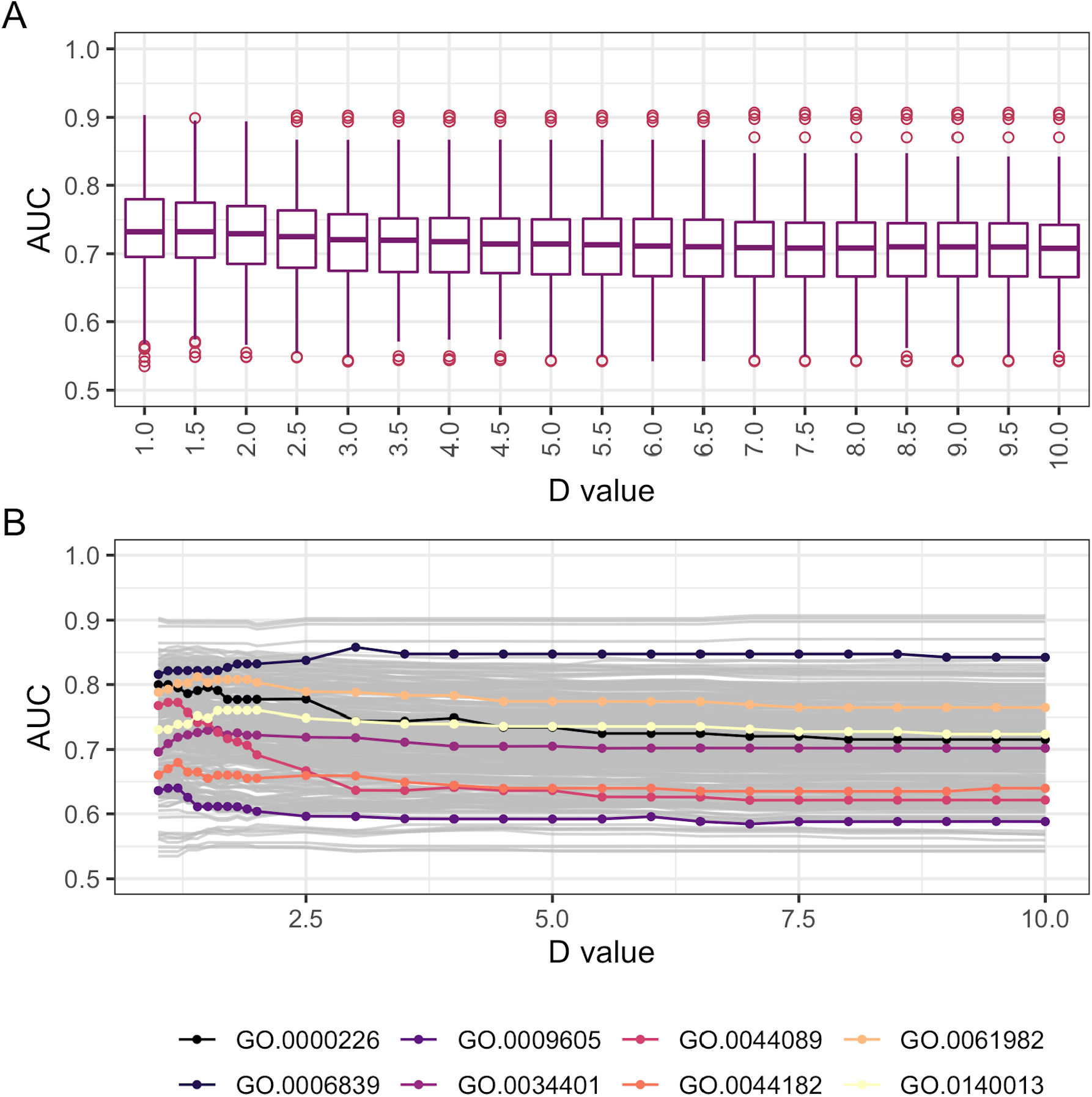
The *D*-value integration parameter. A. Area under curve (AUC) of receiver operator characteristic plots for 298 Gene Ontology biological processes at different *D*-values. B. Variation of AUC for 298 Gene Ontology biological processes at different *D*-values. Eight individual processes are highlighted as exemplars.

## Discussion

Since PFINS are generally created in order to support or derive new biological hypotheses, it is important to generate integrated networks which are as complete and unbiased as possible [22]. Previous PFIN integration techniques have had several drawbacks due to their reliance on an external Gold Standard for dataset scoring. Experimental datasets’ score differently depending upon the Gold Standard chosen [26, 32].

One popular Gold Standard for dataset scoring is the Gene Ontology (GO) [10, 22, 23, 33–35]. The use of GO as a Gold Standard presents some problems due to the hierarchical nature of the database; terms are connected to one another in a parent to child hierarchy, with annotation to a child term automatically implying annotation to all parent terms of that term. Many high level terms, for instance metabolic process (GO:0008152), are too general to imply a realistic functional association [26, 36]. Assuming a functional link based on this term would add noise to the Gold Standard by linking many protein pairs which do not participate in the same cellular process. This problem may be overcome to some extent by ignoring the high-level terms [22, 37]. However, despite GO’s hierarchical nature, the level of a term in the DAG is not necessarily indicative of a term’s specificity [38, 39].

In addition, GO has a number of evidence types with some GO annotations considered more reliable than others [40]. Annotations with the IEA evidence code are considered the least reliable since they are not manually-curated. While the remaining GO annotations are curated, the different evidence types are also thought to differ in their accuracy. In particular, computational evidence codes are generally considered to be less accurate than the experimental codes, and the evidence with the codes inferred from sequence or structural similarity (ISS), inferred from expression pattern (IEP) and non-traceable author statement (NAS) are considered lower reliability than the other codes of their class [41]. Therefore, the choice of what data to include in a GO-based Gold Standard is highly complex and can affect the final network. In our study we chose two different thresholds based on annotation number which resulted in similar number of proteins but vastly different number of interactions in the Gold Standard datasets. While these may not be the optimal datasets derived from GO, it is clear from our results that ssNet produces comparable performance to a GO-derived Gold Standard. In particular, the LTP_GS and BP100_GS-derived networks had the most similar performance with very few statistically significant differences.

Gold Standards from metabolic pathway databases are also commonly used [1, 19, 20]. However, finding a suitable up-to-date metabolic dataset, that has no redundancy with the data to be scored, is hard. Our 2012 study was able to use monthly updates of the KEGG dataset for comparison with monthly updates of BioGRID [8]. However, KEGG is no longer freely available and, although BioSystems represents a comprehensive collection of metabolic pathways that includes KEGG, it has not been updated since 2017.

While it was not possible to find an ideal metabolic Gold Standard for comparison, it is clear from our results that using a dataset with a different focus than the data to be scored, in this case primary metabolic interactions versus experimental interactions, can result in loss of data. Notably, the four LTP yeast data types that did not score against BioSystems (Figure 4 B)—Positive Genetic, Protein-RNA, Synthetic Haploinsufficiency and Affinity Capture-RNA—were genetic/RNA data types. Similarly, many of the HTP datasets lost due to not scoring against BioSystems (Figure 2) were genetic interaction types. Of those lost that were physical interaction data, the vast majority had a non-metabolic focus and four were large-scale *>*2000 interaction datasets [42–45].

If a network study’s focus is non-metabolic, then a metabolic Gold Standard may not be suitable as relevant datasets may be lost. There were far fewer datasets with no score using ssNET and so fewer datasets were discarded. Scoring and integration from a single data source, such as BioGrid, gives a far more complete network without bias towards specific data types. Importantly, since the data can be spilt into individual source studies, this method also ensures no redundancy between data and Gold Standard, which may bias the network.

A metabolic-focused Gold Standard may be desirable if metabolic pathways are the area of interest for downstream analyses, and this is reflected in some of the individual dataset scores. Several datasets scored higher using the metabolic Gold Standard [46–48]. However these were relatively small changes, and while some biological process terms also had better prediction using BioSystems, ssNet had comparable performance for all, in addition to the advantages of the computational ease of using a single source. Furthermore, since key molecular pathway Gold Standards, such as BioSystems, are no longer being maintained/freely-available, at some point they will no longer be sufficiently up-to-date for use as a Gold Standard.

The single source method also overcomes any need for potentially error prone identifier mapping [25]. Identifier mapping required significant effort during creation of our externally-scored networks; although mapping tools were used, some redundancy of identifiers remained that needed time-consuming manual curation. In this case we only counted entrez IDs that could be mapped to gene symbols, however, it should be noted that there were approximately 300 proteins and 1000 interactions that were in the BioSystems dataset and could not be mapped, the majority of which did not overlap with V186 BioGRID data when manually checked. Therefore if ID mapping could have been fully achieved, it would not have improved scoring results. This discrepancy is reflected in the lack of overlap between BioSystems and our source network data.

Yeast remains by far the most complete interaction dataset with data covering almost all of its ∼ 6600 genes^2^[49]. Data coverage in yeast has levelled off since our initial study [8], in particular the fluctuations in protein coverage seen prior to 2010 are no longer observed, reflecting BioGRID’s efforts to improve and standardise curation methods [9, 50]. It is clear that both the data increase and the data curation have affected dataset scores over time. For example, the removal of ubiquitination data from BioGRID [50] resulted in some of the larger changes we observed. Other changes were the direct result of increasing LTP data volume.

Although the overlap in terms of interactions between HTP and LTP data remains relatively low, it still provides enough overlap to allow scoring. While several datasets scored infinity against the BioSystems and GO-derived Gold Standards, very few did so using the LTP data. Infinity scores are hard to deal with since they must be assigned an arbitrary high score (see Methods). However, this is not ideal as demonstrated by the scores of the nine datasets in Figure 2, which have a range of scores using LTP data. By reducing or removing entirely both zero and infinite scoring datasets, an LTP Gold Standard produces more accurate scoring.

The main disadvantage of many PFINs is that the data considered highest quality is used in scoring and not included in the final network [1, 22]. We used an iterative method to score the LTP data by type, before integration with the HTP datasets. While the LTP scores were generally higher than HTP (Figure 4 A), they are in an overlapping range indicating a difference in quality between the data types, and that they can be integrated without introducing significant bias towards the LTP data.

Interestingly, the functional prediction performance of the ssNet networks did not differ from that of the HTP-only networks to any great extent. However, assessing and comparing networks is not straightforward. Here, we measured the quality of networks by prediction of known annotations to produce a numeric measure of network performance for each biological process. While the network performance was not changed for functional prediction, PFIN analyses are often performed in an exploratory manner. The increase connectivity provided by inclusion of the LTP data is likely to be beneficial for other network-based analysis, for instance in clustering and subnetwork analysis [51–53].

We chose thresholds based on our previous studies in yeast[8, 19] but it is possible that different values will be more appropriate in other species, in particular the LTP threshold of 100 interactions. There are also other ways to split the data than by PubMED ID; some studies contain both HTP and LTP data and/or more than one data type. For instance large-scale HTP screens may also include LTP confirmation of some interactions [54] and studies may combine different data types [55]. However, the focus of many studies is in itself something that can be harnessed during integration [19], and we believe keeping the study as a whole dataset and assessing it as a whole is preferable. Finally, a *D*-value of 1.0 was used in our yeast network during integration (see Methods). Higher *D*-values increase the contribution of highest quality datasets to the network, and the optimal *D*-value is dependent on the area of biology being studied (Figure 7). We have therefore provided networks, dataset scores and AUCs, in order to allow selection of the most appropriate data, method and parameters depending on area of study.

## Conclusions

The ssNet method provides a computationally amenable one-step PFIN integration method for functional interaction data which has a number of benefits:

- PFINs are generated from a single data source without using an external Gold Standard.
- The ssNet PFINs are of a quality that is comparable to that of the external Gold Standard PFINs, and exceeds it by some metrics
- Integration is easier and faster, overcoming the challenges of data redundancy, Gold Standard bias and ID mapping.
- The highest quality Gold Standard data can be consistently integrated into the network adding to its quality.

## Methods

### Data sources

BioGRID archive datasets [27] were downloaded for a fifteen year period at six month intervals^3^ from the first available version (V17, 2006) to the current version (V186, 2020). Entrez ID lists for each species were downloaded from the NCBI database^4^ [56] (*S. cerevisiae* 5^*th*^ June 2020), BioSystems Gold Standard data was downloaded from the NCBI’s FTP server^5^ (version 20170421). Annotation information for evaluation was downloaded from the Gene Ontology Resource^6^ [24, 57]: the Gene Ontology obo format (23^*rd*^ March 2020) and annotation mapping files for each species (*S. cerevisiae* 23^*rd*^ March 2020 [58].

### Dataset scoring and integration

The ssNet method involves a five step scoring and integration (Figure 8). BioGRID data were first split by Pubmed ID into HTP and LTP data based on study size threshold (100 throughout unless otherwise stated) and HTP datasets were treated as separate studies. For consistency dataset names are standardised throughtout the results as [Author].[Pubmed ID]. These datasets were then scored using the LTP data as Gold Standard. LTP data were then split into datasets by data type based on the BioGRID experimental code^7^. Each LTP dataset was scored using the remaining LTP data types as Gold Standard. Finally, the HTP and LTP datasets were integrated.

**Figure 8:**
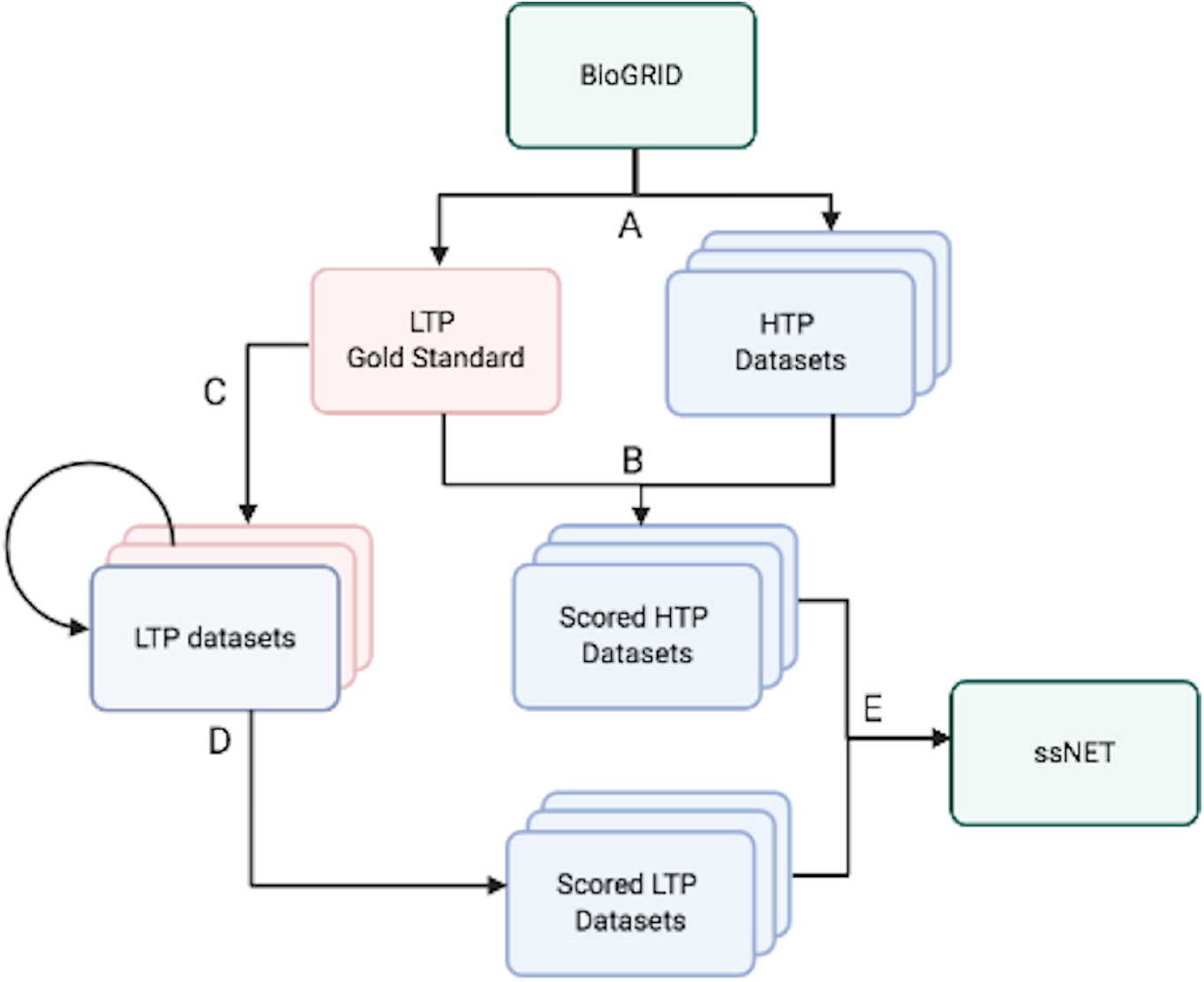
The ssNet scoring and integration method. A. BioGRID data are split into high throughput (HTP) and low throughput (LTP) data based on study size threshold. B. The HTP datasets are scored using the LTP data as Gold Standard. C. LTP data is then split into datasets by experimental type. D. Each LTP data type is scored using the remaining LTP data types as Gold Standard. E. The scored HTP and LTP datasets are then integrated into the final network.

For comparison networks both LTP and HTP datasets were scored against three Gold Standards:

- BioSystems molecular pathway-derived Gold Standard in which proteins in the same pathway were considered Gold Standard pairs (MP_GS).
- A Gene Ontology biological process-derived Gold Standard in which proteins belonging to processes covering <10% of the genome were considered Gold Standard pairs (BP10_GS).
- A Gene Ontology biological process-derived Gold Standard in which proteins belonging to processes with <100 annotation in the yeast genome were considered Gold Standard pairs (BP100_GS).

Confidence scores were calculated using the methods developed by Lee and colleagues [1], which calculates a log-likelihood score for each dataset:

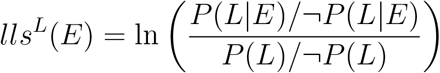

where, *P*(*L*|*E*) and ¬*P*(*L*|*E*) represent the frequencies of linkages *L* observed in a dataset *E* between genes that are linked and not not linked in the Gold Standard, respectively, and, *P*(*L*) and ¬*P*(*L*) represent the prior expectation of linkages between genes that are linked and not not linked in the Gold Standard, respectively. A score greater than zero indicates that the dataset links genes annotated in the Gold Standard. Higher scores indicate greater confidence in the data. Datasets that did not have a positive score or that did not score (*P*(*L*|*E*) = 0) were discarded. Datasets scoring infinity due to perfect overlap with the Gold Standard (¬ *P*(*L*|*E*) = 0) were given a score of ⌈*h*⌉ + 1, where *h* is the set of dataset scores.

Scores were then integrated using the Lee method [1]:

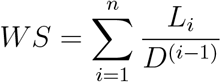

where *L*_1_ is the highest confidence score and *L*_*n*_ the lowest confidence score of a set of *n* datasets. A *D*-value of 1 was chosen for initial network builds, while higher *D*-values were chosen where stated. In HTP-only networks only the scored HTP datasets were integrated.

### Evaluation

Networks were clustered using the Markov Clustering Algorithm (MCL) version 14-137 with default inflation value [31]. Networks were visualised in Cytoscape Version 3.7.2 [59] and the Network Analyser plugin version 3.3.2 was used to calculate topological statistics [60]. Functional prediction was carried out using the Maximum Weight decision rule [61] in which annotations were propagated along the highest weighted edge surrounding a protein. Leave-one-out cross-validation of known annotations was carried out for all GO biological process with at least 100 and no more than 1000 annotations in the genome. Automated annotations with the code Inferred from Electronic Annotation (IEA) were excluded from evaluation. The performance of the networks was evaluated using the area under curve (AUC) of Receiver Operator Characteristic curves for each term [62] using the BioConductor ROC package (v 1.62.0) [63].

The error of the AUC was calculated using the standard error of the Wilcoxon statistic SE(W) [62, 64]:

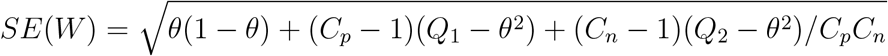

where, *θ* is the AUC, *C*_*p*_ is the number of positive examples, *C*_*n*_ is the number of negative examples and *Q*_1_ and *Q*_2_ are the probabilities of incorrect annotation assignment as defined by:

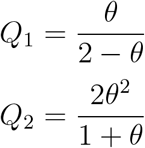

## Competing interests

The authors declare that they have no competing interests.

## Author’s contributions

KJ designed the study and conducted the analyses. KJ and MP wrote the network integration code. KJ, MP, AW and SJC and AA wrote the manuscript. All authors read and approved the final manuscript.

## Footnotes

1 https://thebiogrid.org

2 79% verified; https://www.yeastgenome.org/genomesnapshot on 15th July 2020

3 https://downloads.thebiogrid.org/BioGRID/Release-Archive/

4 https://www.ncbi.nlm.nih.gov/

5 https://ftp.ncbi.nih.gov/pub/biosystems/

6 http://www.geneontology.org/

7 https://wiki.thebiogrid.org/doku.php/experimental_systems

## Notes

https://figshare.com/projects/Yeast_ssNet/114366

https://github.com/kj-intbio/ssnet/

### Competing Interest Statement

The authors have declared no competing interest.

